# Microtubule associated proteins and motors required for ectopic microtubule array formation in *S. cerevisiae*

**DOI:** 10.1101/2021.01.12.426450

**Authors:** Brianna R. King, Janet B. Meehl, Tamira Vojnar, Mark Winey, Eric G. Muller, Trisha N. Davis

## Abstract

The mitotic spindle is resilient to perturbation due to the concerted, and sometimes redundant, action of motors and microtubule-associated proteins. Here we utilize an inducible ectopic microtubule nucleation site in the nucleus of *Saccharomyces cerevisiae* to study three necessary steps in the formation of a bipolar array: the recruitment of the γ-tubulin complex, nucleation and elongation of microtubules, and the organization of microtubules relative to each other. This novel tool, an Spc110 chimera, reveals previously unreported roles of the microtubule-associated proteins Stu2, Bim1, and Bik1, and the motors Vik1 and Kip3. We report that Stu2 and Bim1 are required for nucleation and that Bik1 and Kip3 promote nucleation at the ectopic site. Stu2, Bim1, and Kip3 join their homologs XMAP215, EB1 and kinesin-8 as promoters of microtubule nucleation, while Bik1 promotes MT nucleation indirectly via its role in SPB positioning. Further, we find that the nucleation activity of Stu2 *in vivo* correlates with its polymerase activity *in vitro*. Finally, we provide the first evidence that Vik1, a subunit of Kar3/Vik1 kinesin-14, promotes microtubule minus end focusing at the ectopic site.

## Introduction

High fidelity distribution of genetic material during mitosis requires a dramatic reorganization of microtubules (MTs): the transition from the radiating network of MTs during interphase to the tightly localized bipolar spindle. To form this ordered mitotic array, nucleation of new MTs must be initiated and controlled, and existing MTs must be re-organized.

Successful spindle assembly requires the concerted action of MT organizing centers (MTOCs), motors, and MT associated proteins (MAPs). MTOCs define the distal ends of the spindle and position the spindle in the cell. Motors and MAPs, on the other hand, bind to the MT lattice and have a large variety of functions, from regulating the dynamic instability of MTs to facilitating MT transport.

Several factors complicate our understanding of the roles that MAPs play. First, MAPs act en masse on the multi-micrometer long spindle; many copies of a single protein or a protein complex work together to perform their function efficiently (Page *et al*. 1994; Geiser *et al*. 1997; Miller *et al*. 1998; Miller *et al*. 2000; Derr *et al*. 2012; Furuta *et al*. 2013). Second, MAPs can function either redundantly (Hoyt *et al*. 1992; Roof *et al*. 1992; O’Connell *et al*. 1993; DeZwaan *et al*. 1997; Cottingham *et al*. 1999) or antagonistically (O’Connell *et al*. 1993; Saunders *et al*. 1997). The concerted MAP interactions result in a bipolar spindle that is resilient to perturbation. While this resiliency is advantageous for life, the interplay among MAPs makes assigning roles to individual MAPs difficult. Therefore, we utilized a tool that promotes MT array formation at a site that is ectopic to the bipolar spindle—an inducible Spc110 chimera (Spc110c(T)) (Figure 1). Spc110 is a key component of the yeast spindle pole body (SPB) that recruits the conserved γ-tubulin complex to the poles. Using the chimera-based assay, we have measured the contribution of five highly conserved families of MAPs and motors to MT array nucleation and organization. The tumor-overexpressed gene (TOG) polymerase Stu2, the end-binding (EB) protein Bim1, the CLIP170 homolog Bik1, and the kinesin-8 and kinesin-14 motor proteins, Kip3 and Vik1, were examined. Each have been previously implicated in MT nucleation and spindle assembly.

**Figure 1.**
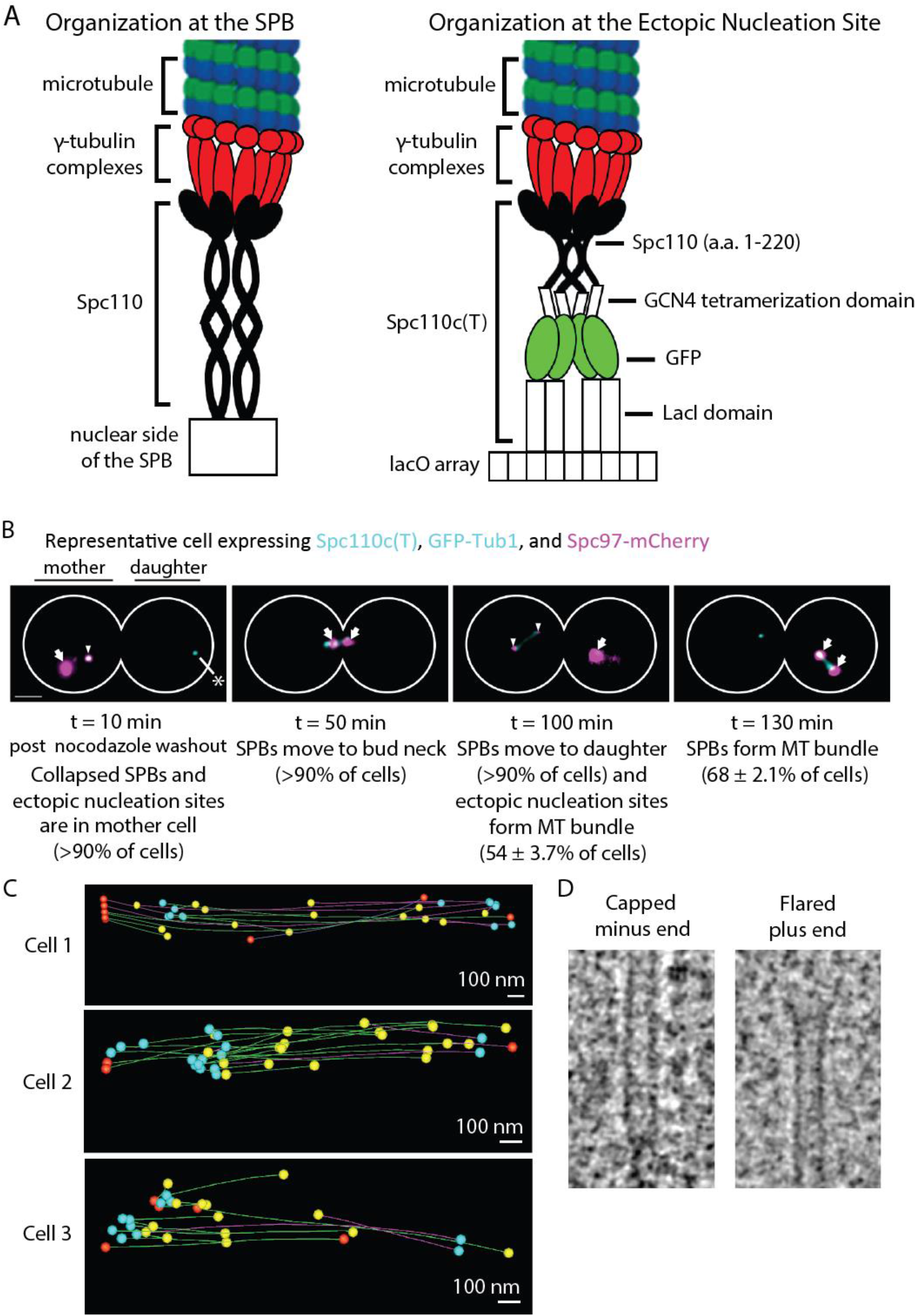
The γ-tubulin complex serves as a robust MT nucleation template at the wild-type SPB and the Spc110c(T). (A) (Left) Full-length Spc110 dimerizes via a coiled coil domain and anchors γ-tubulin complexes to the nuclear side of the budding yeast SPB. (Right) The Spc110c(T) is comprised of the N-terminal 220 amino acids of Spc110, a tetramerization version of the GCN4 domain, a GFP tag, and a lac repressor domain which binds *lacO* arrays integrated into Chromosome XII. (B) A representative timeline of events following nocodazole arrest, β-estradiol pulse induction of Spc110c(T), and nocodazole release (RKY52). At the earliest timepoints, SPBs are collapsed and the Spc110c(T) is colocalized with Spc97-mCherry. SPBs then move to the bud neck and into the daughter cell in 92 ± 4.6% of cells. Following SPB movement to daughter, the Spc110c(T) forms MT arrays in the mother and the SPBs form spindles. γ-tubulin complexes are visualized using expression of Spc97-mCherry (shown in magenta), and αβ-tubulin is visualized using expression of GFP-Tub1 (shown in cyan). Arrows denote SPBs and carets denote ectopically localized γ-tubulin complexes. Asterisk in left panel denotes a commonly observed phenomenon following Spc110c(T) expression—large GFP puncta in daughter cell that do not recruit Spc97-mCherry. Scale bar is 2 μM. (C) Electron tomography of MT bundles associated with Spc110c(T) revealed antiparallel MTs with clusters of capped minus ends (TVY11-123D). Capped minus ends shown in blue, flared plus ends shown in yellow, and MT ends out of volume shown in orange. Green MTs have minus end left of their plus end, and pink MTs have opposite polarity. Polarity of MTs was assessed by their visible ends. (D) Representative capped minus end and flared plus end from electron tomography.

*Saccharomyces cerevisae* has a closed mitosis, and nucleation of new MTs during spindle assembly is narrowly localized to the nuclear face of the SPB. Of the five candidate budding yeast proteins, four have homologs that are implicated in MT nucleation in other organisms. Though these proteins primarily bind the MT lattice, four interact with components of the SPB: Stu2, Bim1, Bik1, and Vik1 (Wigge *et al*. 1998; Manning *et al*. 1999; Newman *et al*. 2000; Usui *et al*. 2003; Wang *et al*. 2012). Furthermore, homologs of Stu2, Bim1, and Kip3 were shown to promote MT nucleation both *in vivo* and *in vitro* (Popov *et al*. 2002; Vitre *et al*. 2008; West and McIntosh 2008; Erent *et al*. 2012; Wieczorek *et al*. 2015; Thawani *et al*. 2018), and Stu2 specifically promotes cytoplasmic MT nucleation (Gunzelmann *et al*. 2018). Conversely, Plk1, the fission yeast kinesin-14 motor, suppresses MT nucleation (Olmsted *et al*. 2014).

Spindle assembly also requires careful regulation of dynamic instability at the MT plus ends, which extend away from the SPB and connect to the kinetochores. In the case of Stu2, Bim1, and Bik1, there is evidence of MT stabilization for each. Stu2 is a member of the XMAP215 family of MT polymerases that promote growth rates (Brouhard *et al*. 2008). Bim1 inhibits catastrophe and does so in complex with Bik1 (Blake-Hodek *et al*. 2010). Kip3, on the other hand, destabilizes long MTs and stabilizes short MTs (Gupta *et al*. 2006; Varga *et al*. 2006; Tischer *et al*. 2009; Gardner *et al*. 2011; Reese *et al*. 2011; Su *et al*. 2011; Melbinger *et al*. 2012; Fukuda *et al*. 2014).

In addition to regulating the length of MTs, MAPs or motors can function to organize MTs relative to each other. Specifically, interpolar MTs are organized to overlap in the midzone, while kinetochore MTs are focused toward MTOCs via their minus ends in order to generate tension. Dynein and kinesin-14 motors promote MT minus end focusing at the MTOCs in *D. melanogaster* and *H. sapiens* (Endow and Komma 1998; Kwon *et al*. 2008; Lecland and Lüders 2014). No such activity has been reported in budding yeast; however mutations that weaken the nuclear SPB-MT interface are synthetically lethal with Kar3 and Vik1 (of the Kar3Vik1 complex) (Greenland *et al*. 2010).

## Materials and Methods

### Media

YPD was prepared as described (Burke *et al*. 2000). YPD 3x ADE media is YPD with 5 mg/ml adenine. LoFlo S liquid medium contains 1.93 g/l yeast nitrogen base LoFlo (Formedium^™^) plus 5 g/l ammonium sulfate. LoFlo SD^+^ medium is LoFlo S medium with 2% glucose and 1/100 dilution of a filter sterilized solution of L-amino acids and nutrients containing 0.2% arginine, histidine and methionine, 0.3% isoleucine, leucine, and lysine, 0.4% tryptophan, 0.5% phenylalanine and threonine, 0.25% uracil and 1.5% adenine. The high adenine suppressed the formation of red fluorescent pigment in our *ade2, ADE3* strains.

### Strain construction

All strains used in this study are derivatives of W303 (Thomas and Rothstein 1989) (Table S1). Transformations were performed by a standard lithium acetate method (Gietz *et al*. 1995). The tetrameric Spc110 chimera construct, Spc110c(T), consisting of *SPC110(1-220a*.*a*.*)-GCN4-GFP-lacI* was integrated at *ADE2* with a promoter specific to the transcription factor Z_4_EV as previously described (Lyon et al., 2016). Z_4_EV was integrated at *CAN1* with a β-estradiol inducible promoter (McIsaac et al., 2013). 256 copies of a lacO repeat were integrated into chromosome XII as described previously (Lyon et al., 2016). MAPs were C-terminally tagged with 3 V5 epitopes and the auxin-responsive protein IAA7 (Shetty *et al*. 2019). The ubiquitin ligase complex protein OsTIR1, necessary for auxin-inducible degradation, was integrated at *LEU2*. Spc97 was C-terminally tagged with mCherry. GFP-Tub1 under the *TUB1* promoter was integrated at URA3 using pAFS125 (Straight *et al*. 1997).

### Electron tomography

TVY11-123D was incubated with 1 mM auxin (Millipore Sigma, St. Louis, MO) for 2.5 hours (Spc110-AID degradation greater than 90%). Cells were collected on a Millipore filter by vacuum filtration and high pressure frozen using a Wohlwend Compact 02 high pressure freezer (Technotrade International, Manchester, New Hampshire). Cells were freeze substituted at low temperature in acetone containing 0.25% glutaraldehyde and 0.1% uranyl acetate, then infiltrated with Lowicryl HM20 resin. After polymerization, 350 nm serial sections were collected on copper slot grids coated with formvar, post-stained with 2% aqueous uranyl acetate and Reynold’s lead citrate. Tilt series were collected on a Technai F30 microscope. Tomograms were computed and modeled using the IMOD software package (Kremer *et al*. 1996; Mastronarde 1997; O’Toole *et al*. 2002).

### γ-tubulin complex recruitment assay

Cells were grown asynchronously in YPD 3xADE at 25°C to 40 Klett units, within log phase. Cells were first pulsed with 0.1 μM β-estradiol (Millipore Sigma) for 10 minutes and then washed with YPD 3xADE. Cells were then treated with 15 μg/mL nocodazole (Millipore Sigma) for 2.5 hours. In cells harboring a protein tagged with auxin-inducible degron, cells were simultaneously treated with 1 mM auxin (Millipore Sigma) during the hour incubation. Cells were pelleted and resuspended in 100 µl of LoFlo SD medium, placed on Lo-Flo SD + 1% agarose pad (SeaKem ® Gold Agarose, Lonza, Rockland, ME) supplemented with 15 μg/mL nocodazole and 1 mM auxin, as previously described (Muller et al., 2005) https://www.youtube.com/watch?v=ZrZVbFg9NE8).> Cells were imaged using either a DeltaVision system (Applied Precision, Issaquah, WA) or an Axio Observer (Carl Zeiss). The DeltaVision system was equipped with IX70 inverted microscope (Olympus, Center Valley, PA), a Coolsnap HQ digital camera (Photometrics, Tucson, AZ), and a U Plan Apo 100X objective (1.35 NA). The inverted Axio Observer was equipped with an ORCA-Flash4.0 V3 Digital CMOS camera (Hamamatsu), a Spectra X LED light engine (Lumencor), a Plan-APOCHROMAT 63X/1.46 objective (Zeiss), and a 432/523/702 nm BrightLine® triple-band bandpass filter (Semrock, Inc., Rochester, NY).

For the recruitment assay, the Spc110c(T) was visualized using its integrated GFP fluorescence and γ-tubulin complexes were visualized with Spc97-mCherry. GFP was imaged with 0.25 s exposures, mCherry was imaged with 0.4 s exposures, and 25 0.2 µm z-sections were taken. Images taken on the DeltaVision microscope were binned 2 × 2, while images taken on the Axio Observer were not binned.

Recruitment analysis was performed using the Spots colocalization function of Imaris software (Bitplane, Switzerland) (RRID: SCR_007370). Spc110(1-220a.a.)-GCN4-GFP-lacI puncta were located by assigning an XY diameter of 0.4 μm and a Z diameter of 1.3 μm. Spc97-mCherry puncta were located by assigning an XY diameter of 0.4 μm and a Z diameter of 1.8 μm. Quality of spots was manually determined. To exclude background fluorescence of the agarose pad in the total number of puncta, GFP and mCherry fluorescent puncta were first colocalized to be within 4 μm. These filtered spots were then colocalized within 0.5 μm and reported as the percent of Spc110c(T) puncta with recruitment events.

### MT formation assay

Cells were grown asynchronously in YPD 3xADE at 25°C to 40 Klett units, within log phase. Cells were then treated with 15 μg/mL nocodazole for 1 hour. 1 mM auxin was then added, and the incubation continued for an additional hour. 0.1 μM β-estradiol was then added, and the incubation continued for an additional 1.5 hours. Cells were released from nocodazole and β-estradiol and treated with only 1 mM auxin for 10 minutes. Cells were pelleted and placed on Lo-Flo agarose pads supplemented with auxin. Cells were imaged at 10-minute intervals for 2.5 hours as described above for the γ-tubulin recruitment assay.

For the MT formation assay, the Spc110c(T) and Spc97-mCherry were visualized and acquired as described for the recruitment assay, with MTs visualized using GFP-Tub1 and only 15 0.2 µm z-sections.

MT formation analysis was performed manually using Imaris software (Bitplane).

### Calculations for an indirect effect of Bik1

In control cells where SPBs move to the daughter, the fraction of those cells that form Spc110c(T)-associated MT arrays, or A_daughter_, is 0.58. When SPBs remain in the mother, the fraction, or A_mother_, is 0.083. When Bik1-AID is degraded, the fraction of cells with SPBs remaining in the mother, or SPB_mother_, is 0.95, while the fraction of cells with SPBs moving to the daughter, or SPB_daughter_, is 0.048. If the effect of Bik1-AID on ectopic MT array formation is due purely to SPB localization, the predicted fraction of cells with Spc110c(T)-associated MT arrays is 0.11, as given by the following equation:

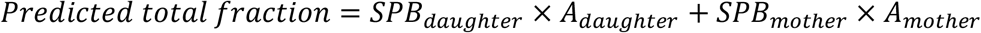

### Line scan analysis

Ten-pixel wide lines along Spc110c(T)-associated MT arrays were drawn manually using ImageJ (National Institutes of Health, United States), and measured values were the average fluorescence intensity at each point on the line scan. Background fluorescence was calculated using the average of the last five points of the line scan. After subtraction of background, line scans were analyzed using the peakfinder and trapz functions of Matlab (MathWorks, United States).

## Results and Discussion

### Spc110c(T) recruits the γ-tubulin complex and forms MT arrays *in vivo*

Temporally controlled by an inducible promoter, the Spc110c(T) recruits the γ-tubulin complex to the DNA, rather than the SPB, through lac operator/lac repressor interactions (Figure 1). This provides an ectopic site of nucleation that is distinct from the SPB. Over a three-hour observation period, 68 ± 2.1% cells form MT arrays at the SPB, while 54 ± 3.7% of cells form ectopic MT arrays at the Spc110c(T). Arrays at the Spc110c(T) are composed of antiparallel MTs with capped minus ends, as revealed by electron tomography (Figure 1C). While Spc110c(T)-associated MT bundles are less organized than mitotic spindles, MT minus ends cluster.

### MAPs and motors are not required for localization of γ-tubulin complex to the Spc110c(T)

An initial, necessary step in MT nucleation is the recruitment of the γ-tubulin complex to Spc110 at the SPB. When cellular MTs are depolymerized in 15 μM nocodazole, the Spc110c(T) recruits the γ-tubulin complex, which is quantified as the percent of Spc110c(T) puncta that ectopically colocalize with Spc97-mCherry puncta (Figure 2). As described above, several of the candidate MAPs and motors interact with components of the SPB, so we asked whether the candidate proteins affected the ability of the N-terminus of Spc110 to recruit or bind γ-tubulin complexes. We tested the effects of deletion of *VIK1* and *KIP3* and the effect of depletion of Stu2, Bim1, and Bik1 using an auxin-degradation system. We found that none of the MAPs or motors promote recruitment of the γ-tubulin complex to Spc110c(T) (Figure 2).

**Figure 2.**
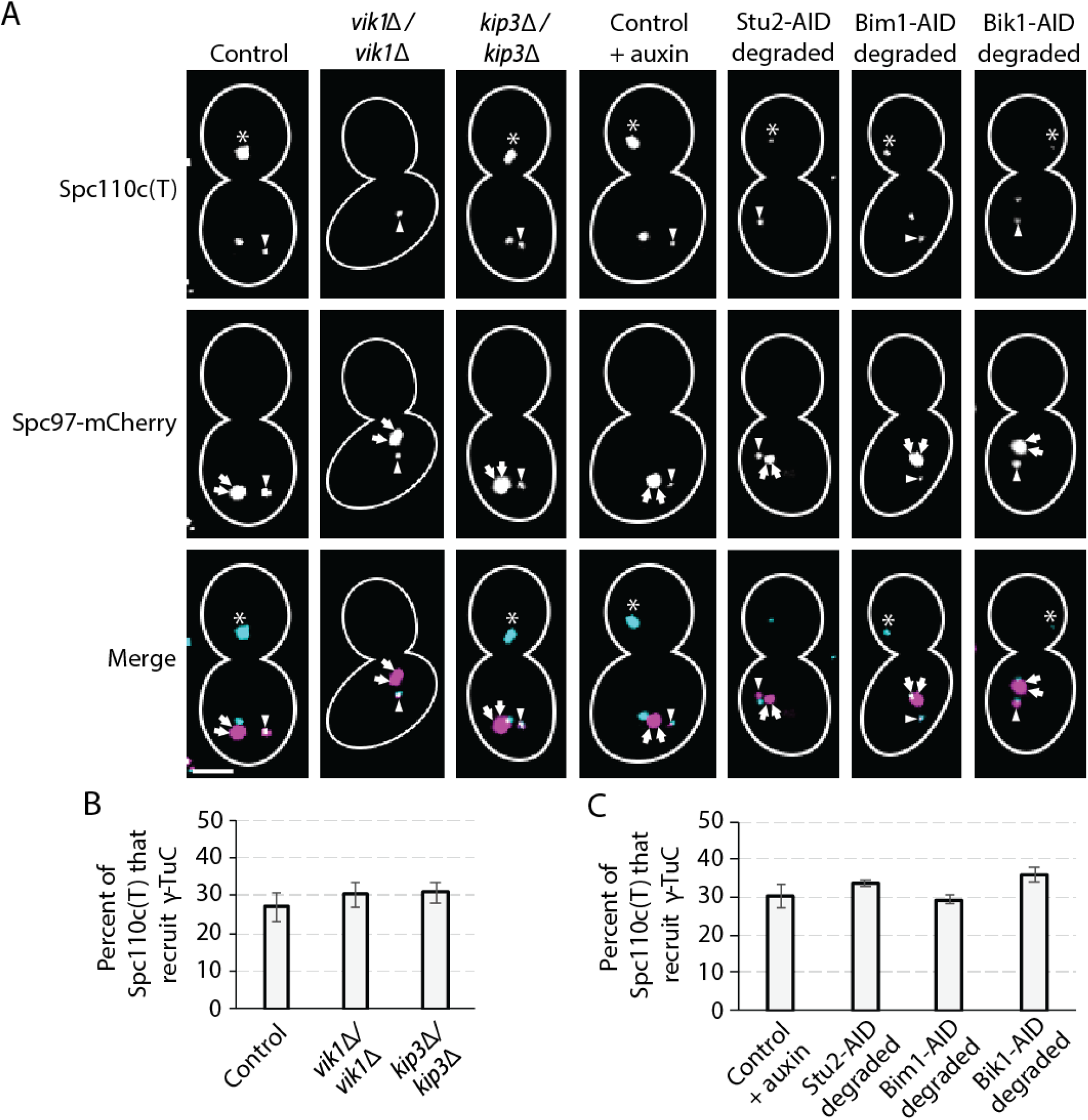
MAPs and motors are not required for γ-tubulin complex localization to the Spc110c(T). (A) Representative recruitment of γ-tubulin complex to Spc110c(T) in control (RKY44), *vik1Δ/vik1Δ* (RKY174), *kip3Δ/kip3Δ* (RKY56), and auxin-treated control, Stu2-AID (RKY14), Bim1-AID (RKY47), and Bik1-AID (RKY51) cells. γ-tubulin complexes are visualized using expression of Spc97-mCherry (shown in magenta). Arrows denote SPBs and carets denote ectopically localized γ-tubulin complexes. Asterisks denote GFP puncta in daughter cell that do not recruit Spc97-mCherry and are in the cytoplasm. Scale bar is 3 μM. (B) Percent of cells in which the Spc110c(T) recruits the γ-tubulin complex is unaffected by deletion of Vik1 or Kip3. Values shown are mean ± S.D. There were 2 biological replicates for all conditions and 234, 316, and 375 total cells for control, *vik1Δ/vik1Δ*‚and *kip3Δ/kip3Δ*‚ respectively. (C) Percent of cells in which the Spc110c(T) recruits the γ-tubulin complex is unaffected by degradation of Stu2, Bim1, or Bik1. Values shown are mean ± S.D. There were 2 biological replicates for all conditions and 249, 389, 248, and 315 total cells for control plus auxin, Stu2-AID degraded, Bim1-AID degraded, and Bik1-AID degraded, respectively.

### Stu2, Bim1, Bik1, and Kip3 promote MT formation at the Spc110c(T)

Following nocodazole treatment and subsequent washout, the Spc110c(T) promotes formation of ectopic MT arrays in 54 ± 3.7% of cells, (32 ± 1.5% with auxin treatment) (Figure 3). We tested the role of Vik1, Kip3, Stu2, Bim1, and Bik1 in this phenomenon by quantifying the cumulative percent of cells forming MT arrays at both the SPBs and ectopic sites. We found that Vik1 was not required for MT array formation at either the SPB or the Spc110c(T) (Figures 3B and 3C). Strikingly, Stu2 and Bim1 are essential for MT formation from both the Spc110c(T) and the SPB. No visible MTs form when either Stu2-AID or Bim1-AID is degraded (78% and 99% degradation, respectively) (Figures 3B and 3C).

**Figure 3.**
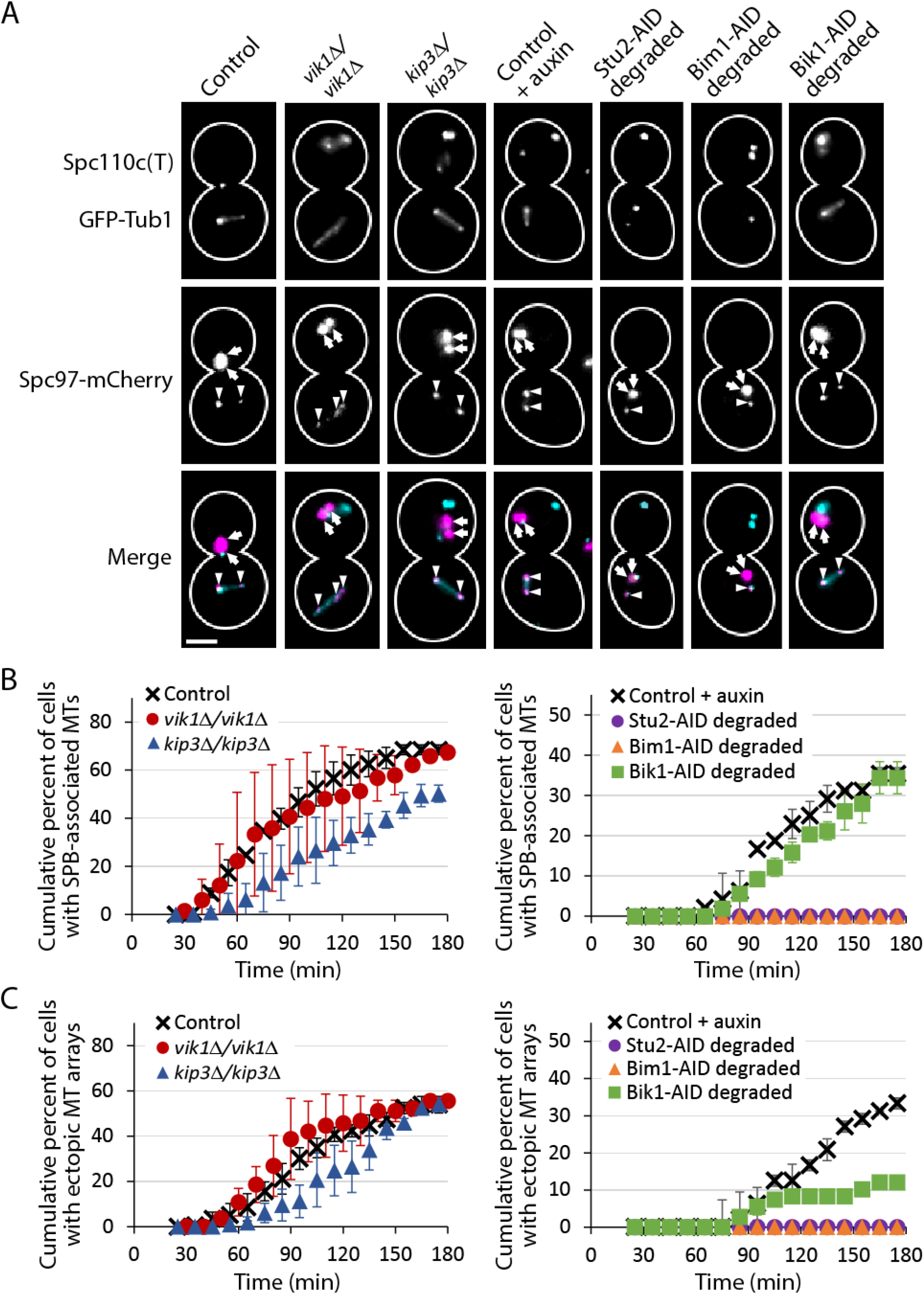
Stu2, Bim1, Bik1, and Kip3 promote MT array formation from the Spc110c(T). (A) Representative MT array formation from the Spc110c(T) in control (RKY52), *vik1Δ/vik1Δ* (RKY31), *kip3Δ/kip3Δ* (RKY49), and auxin-treated control, Stu2-AID (RKY7), Bim1-AID (RKY46), and Bik1-AID (RKY36) strains. γ-tubulin complexes are visualized using expression of Spc97-mCherry (shown in magenta) and αβ-tubulin is visualized using expression of GFP-Tub1 (shown in cyan). Arrows denote SPBs and carets denote ectopically localized γ-tubulin complexes. Asterisks denote GFP puncta in daughter cell that do not recruit Spc97-mCherry. Scale bar is 3 μM. (B) Cumulative percent of cells over time in which SPBs form spindles. Values shown are mean ± S.D. There were ≥ 2 biological replicates and 157, 108, and 120 total cells for control, *vik1Δ/vik1Δ*‚ and *kip3Δ/kip3Δ*‚ respectively. (C) Cumulative percent of cells over time in which Spc110c(T)s form MT arrays. Values shown are mean ± S.D. There were 2 biological replicates for all conditions and 204, 173, 155, and 108 total cells for control plus auxin, Stu2-AID degraded, Bim1-AID degraded, and Bik1-AID degraded, respectively

Although not essential for MT array formation, both Kip3 and Bik1 each promote MT array formation at the Spc110c(T). For Kip3, MT formation is slightly delayed in *kip3Δ/kip3Δ* cells at both the ectopic site and the SPB (Figure 3C).

When Bik1-AID is degraded (82% degradation), MT formation is also delayed at the ectopic site, but by a unique mechanism. While degradation of Bik1-AID does not impact MT array formation between SPBs (Figure 3B), it does largely disrupt the movement of SPBs to the daughter (Figure 4). As seen in control cells, this movement of the SPB to the daughter enhances the formation of the ectopic MT bundle in the mother from 8.3% to 58% (Figure 4A). If the impact of Bik1 degradation on ectopic MT array formation is due to SPB localization, we would expect ectopic MT array formation to be approximately 11% (calculations shown in Materials and Methods). As the observed value is 12 ± 0.36%, the reduction of ectopic MT formation following Bik-AID degradation is most likely indirect and largely a result of the effect on SPB positioning. A role for Bik1 in SPB positioning has previously been observed (Carvalho *et al*. 2004; Caudron *et al*. 2008; McNally 2013).

**Figure 4.**
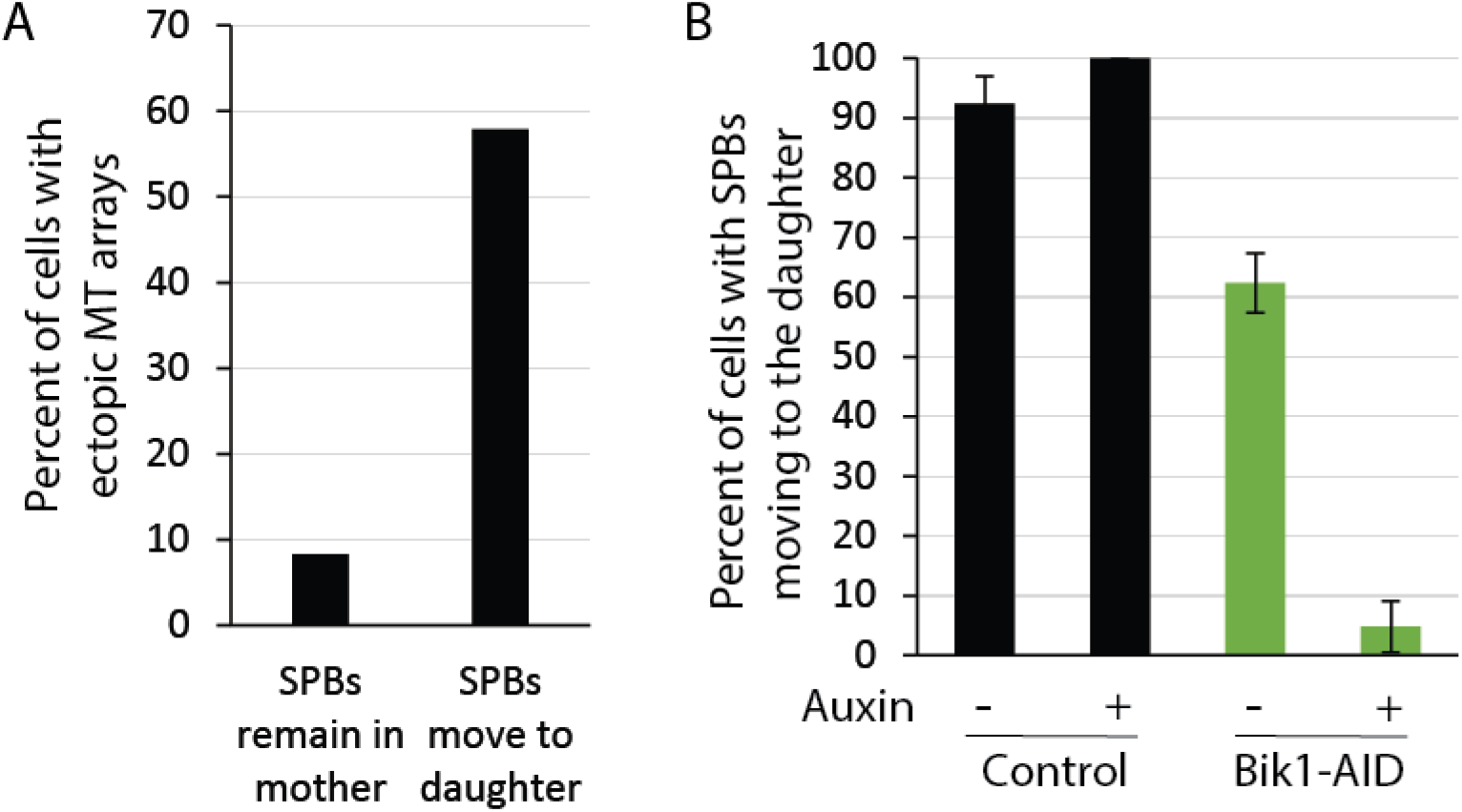
Bik1 indirectly promotes MT nucleation at the Spc110 chimera. (A) SPB movement to daughter cells correlates positively with Spc110c(T) MT formation. During the examination of ectopic MT formation in RKY52, cells were scored for the position of the SPB in the budded cells. (B) Expression of Bik1-AID and depletion of Bik1-AID (RKY36) results in decreased SPB movement to daughter cells. Values shown are mean ± S.D. There were ≥ 2 biological replicates and 157, 204, 100, and 108 total cells, for control, control plus auxin, Bik1-AID, and Bik1-AID plus auxin, respectively.

### The ability of the Spc110c(T) to form MT arrays correlates with the polymerase activity of Stu2

Recent work from our lab showed that the MT polymerization rate *in vitro* of XMAP215, the *X. laevis* homolog of Stu2, correlates with its ability to nucleate MTs in *vitro*. We asked whether previously published MT polymerization rates of Stu2 mutants *in vitro* would correlate with their promotion of MT array formation at the Spc110c(T).

Stu2 polymerase activity can be modulated by inhibiting either its ability to bind free tubulin via TOG domains, to localize to the MT lattice via the basic MT binding domain, or to dimerize via the coiled coil domain. One study has quantified the polymerase activity of various Stu2 mutants, placing them into three categories: wild-type polymerase activity, moderate polymerase activity, or no activity (Figure 5A, data from (Geyer *et al*. 2018).

**Figure 5.**
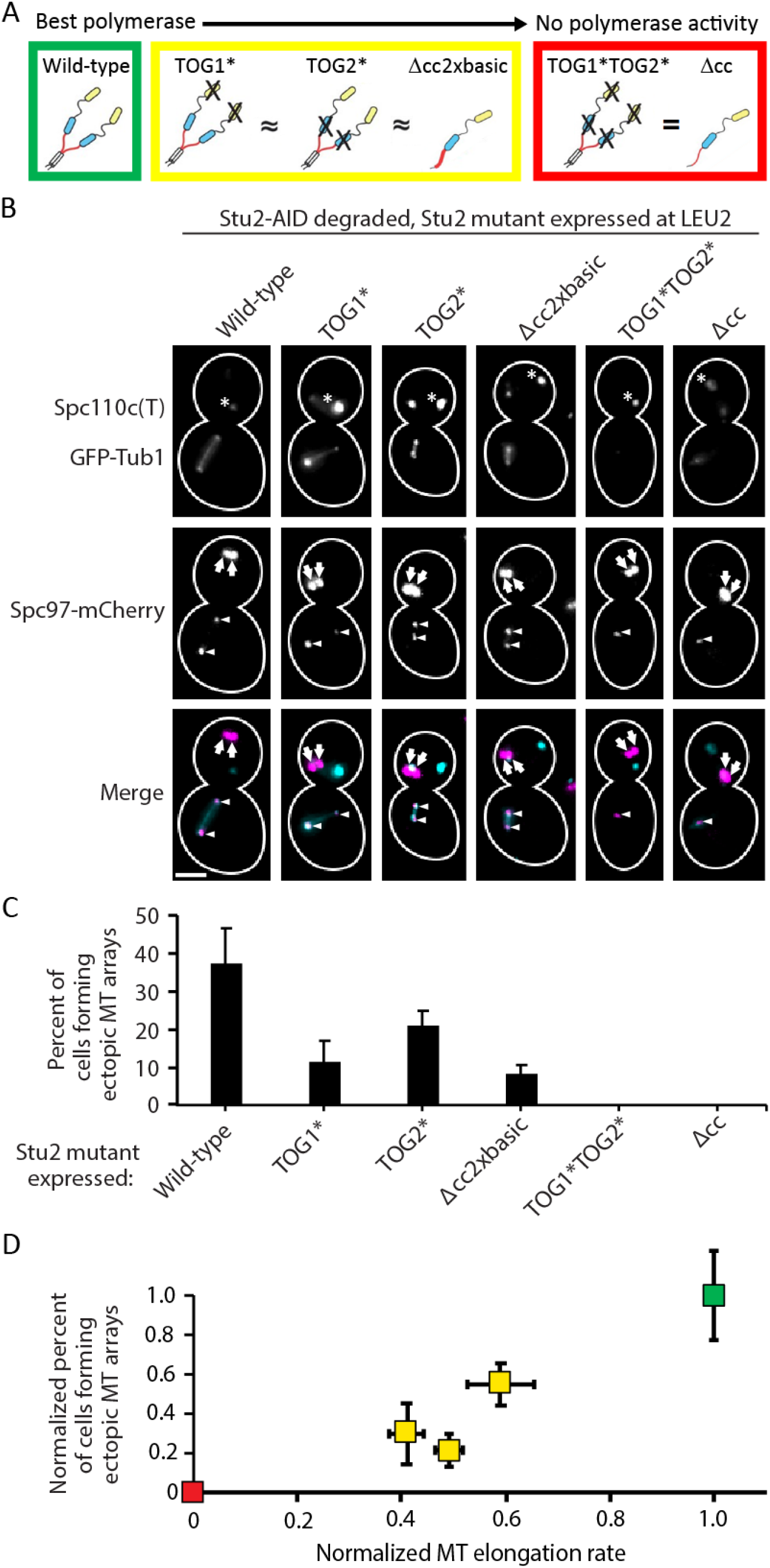
Stu2 promotion of MT nucleation at the Spc110c(T) correlates with its reported polymerase activity *in vitro*. (A) Stu2 mutants used in this study fall into three distinct categories of polymerase activity, as quantified by (Geyer *et al*. 2018). (B) Representative MT array formation at the Spc110c(T) when wild-type Stu2 (RKY122), Stu2 TOG1* (RKY126), Stu2 TOG2* (RKY133), Stu2 Δcc 2xbasic (RKY140), Stu2 TOG1* TOG2* (RKY132), and Stu2 Δcc (RKY139) are expressed. γ-tubulin complexes are visualized with Spc97-mCherry (shown in magenta) and tubulin is visualized with GFP-Tub1 (shown in cyan). Arrows denote SPBs and carets denote ectopically localized γ-tubulin complexes. (C) Percent of cells in which Spc110c(T) forms a MT array over a three-hour period following nocodazole washout. Values are mean ± S.D. There were 2 biological replicates for all conditions and 101, 135, 157, 71, 138, and 154 total cells for Stu2, Stu2 TOG1*, Stu2 TOG2*, Stu2 Δcc 2xbasic, Stu2 TOG1* TOG2*, and Stu2 Δcc, respectively. (D) Percent of cells in which Spc110c(T) forms a MT array plotted against MT elongation rate, where each is normalized to the activity of wild-type Stu2. Data represents the results of Figure 5C and the published results of (Geyer *et al*. 2018). Colors correspond to color of boxes surrounding the depictions of the three classes of mutants in (A).

As shown above, Stu2 is required for MT formation following nocodazole treatment (Figure 3). When Stu2-AID is degraded, MT array formation is rescued through constitutive expression of Stu2 at an exogenous locus (Figure 5). We utilized this system to compare Stu2 mutants with different polymerase activities. Stu2-AID at its native locus was degraded during Spc110c(T) expression, and Stu2 mutants were expressed constitutively at LEU2. As hypothesized, Stu2 polymerase activity *in vitro* correlated with MT formation activity (Figures 5C and 5D).

### Kinesin-14 motor complex protein Vik1 promotes focusing of MT minus ends

While Vik1 was not required for ectopic recruitment of the γ-tubulin complex to Spc110c(T), nor ectopic MT array formation, deletion of *VIK1* did result in decreased MT minus end clustering within the ectopic microtubule array (Figure 6). We quantified this phenotype by measuring Spc97-mCherry intensity along Spc110c(T)-associated MT arrays (Figure 6B). Importantly, the total Spc97-mCherry intensity along Spc110c(T)-associated MT arrays is the same in control cells as in *vik1Δ/vik1Δ* cells (Figure 6C). However, in control cells, Spc97-mCherry usually localizes to two bright puncta at the end of ectopic MT arrays (73% of cells, Figure 6D). In contrast, Spc97-mCherry is more evenly distributed along Spc110c(T)-associated MT arrays in *vik1Δ/vik1Δ* cells. This phenotype was quantified using the root mean squared error (RMSE) (Equation 1):

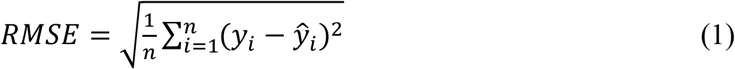

where *n* = number of Spc110c(T) MT bundles measured, *y_i_* = Spc97-mCherry intensity at a given point within the bundle, and *ŷ_i_* = mean Spc97-mCherry intensity across the MT bundles. When γ-tubulin complexes cluster, the Spc97-mCherry intensity at any given point will be brighter and thus further from the mean of the Spc97-mCherry intensity along the MT bundle. Using this measure, Vik1 significantly promotes MT minus end clustering (p <0.001, Figure 6E).

**Figure 6.**
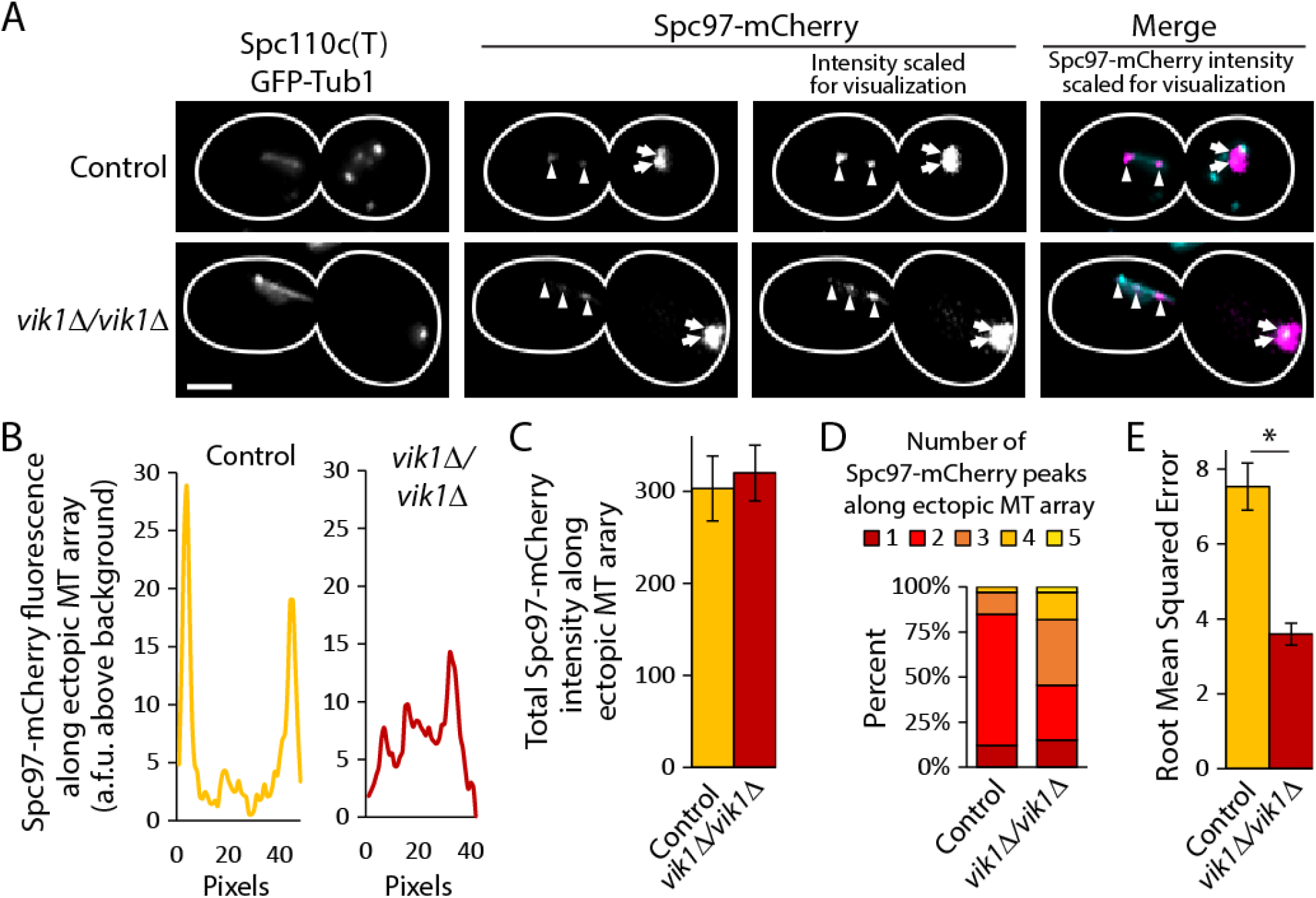
Vik1 significantly promotes γ-tubulin complex clustering. (A) Representative control (RKY52) shows that the Spc110c(T) characteristically forms MT bundles with distinct γ-tubulin complex puncta at each end (top), and in *Δvik1/Δvik1* (RKY31), the Spc110c(T) forms MT arrays with disperse γ-tubulin complexes along the array. (B) Representative line scan of the average Spc97-mCherry intensity at a given pixel along the length of a 10-pixel wide line scan of the Spc110c(T) associated MT array in control (left) and *Δvik1/Δvik1* cells (right). Data plotted is the sum intensity above background fluorescence across a 10-pixel line. (C) Average total Spc97-mCherry intensity along Spc110c(T)-associated MT arrays. (D) Number of Spc97-mCherry peaks along ectopic MT arrays differs between control cells and *Δvik1/Δvik1* cells. (E) Average RMSE of Spc97-mCherry distribution along line scans of Spc110c(T)-associated MT bundles. Data plotted are means ± S.D. (control N = 28, *vik1Δ* N = 33). Asterisk denotes significance as measured by t-test, p<0.001.

The Spc110c(T) provides a unique system to analyze the roles of MAPs and motors whose roles have been obscured in the context of the resilient bipolar spindle. Here we report previously undetected roles for Stu2, Bim1, Bik1, Kip3 and Vik1 in the formation of a bipolar MT array in the nucleus.

The formation of a bipolar array at the Spc110c(T) can be divided into a number of steps, beginning with the recruitment of the γ-tubulin complex to the Spc110c(T). Of the MAPs and motors included in this study, we found that none affected the ability of the Spc110c(T) to recruit the γ-tubulin complex. The ability of the Spc110c(T) to recruit might be strictly due to upstream regulation, such as post-translational modifications and/or oligomerization states of the nucleation machinery (Lyon *et al*. 2016).

The Spc110c(T) also revealed previously unreported roles in MT nucleation for Stu2, Bim1, Bik1, and Kip3 in budding yeast nuclei. Bim1 and Kip3 join their homologs as promoters of MT nucleation (Vitre *et al*. 2008; West and McIntosh 2008; Erent *et al*. 2012). Bik1 becomes the first CLIP170 homolog to be implicated in MT nucleation, but due to its role in SPB positioning, the delay in MT formation at the Spc110c(T) may be an indirect consequence of SPB localization. Stu2, which localizes to the nucleus and the cytoplasm, has now been shown to promote MT nucleation in both compartments (Gunzelmann *et al*. 2018). Further, the polymerase activity of Stu2 *in vitro* correlates well with its nucleation activity *in vivo*, which supports the theory that Stu2 homologs function to elongate nascent MTs during nucleation (King *et al*. 2020).

Finally, we report a role for Vik1 in MT minus end focusing. Kinesin-14 motor proteins have been implicated in spindle repair via MT minus end focusing in other eukaryotic cells (Endow and Komma 1998; Kwon *et al*. 2008; Lecland and Lüders 2014). However, this is the first evidence of this role for Vik1, suggesting a broadly conserved role for the kinesin-14 motor proteins.

## Acknowledgments

We thank Matthew Miller for the plasmids to integrate Stu2 mutants. This study was supported by grants from the National Institutes of Health grant P01 GM105537 to Mark Winey, Trisha Davis, R35 GM130293 to Trisha Davis, and T32 GM007270 to Brianna Rose King.

## Author contributions

BRK designed and performed research, analyzed the data, and wrote the manuscript. JBM performed tomography experiments. TV performed research. MW contributed experimental design, supervised research and edited the manuscript. EGM contributed experimental design and edited the manuscript. TND contributed experimental design, analyzed data, supervised the research, and helped write the manuscript.

## Tables

**Table S1.**
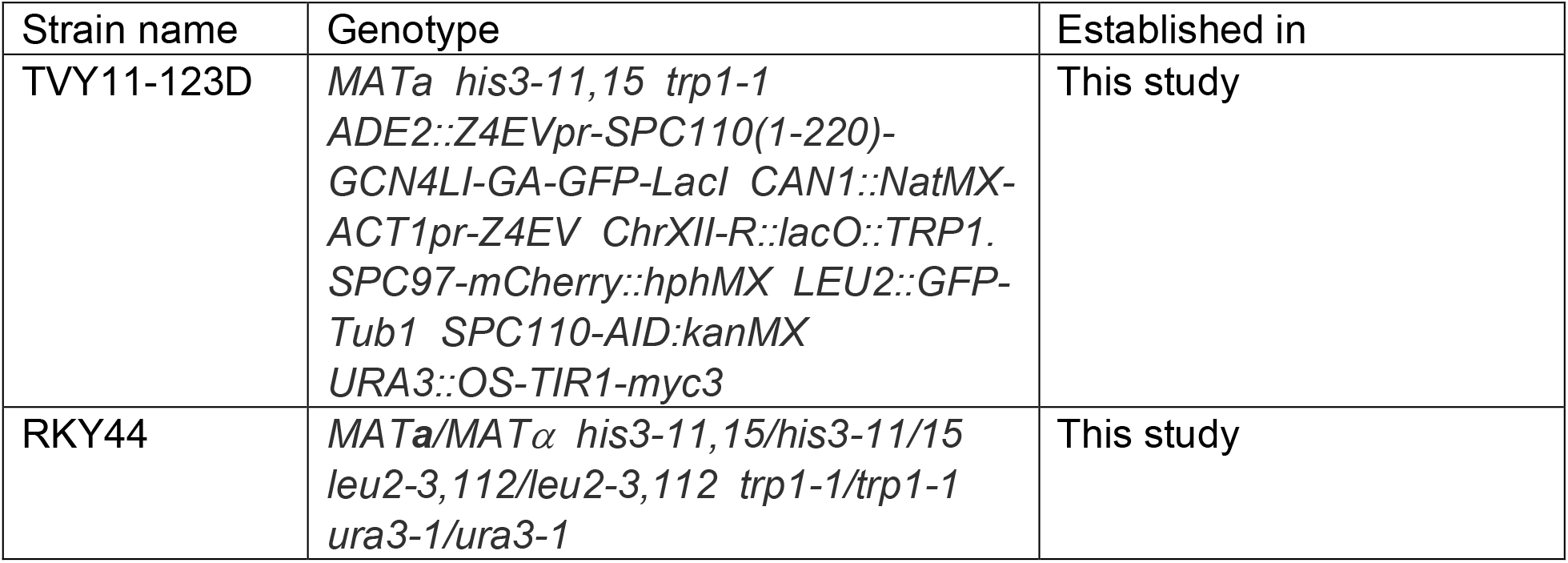
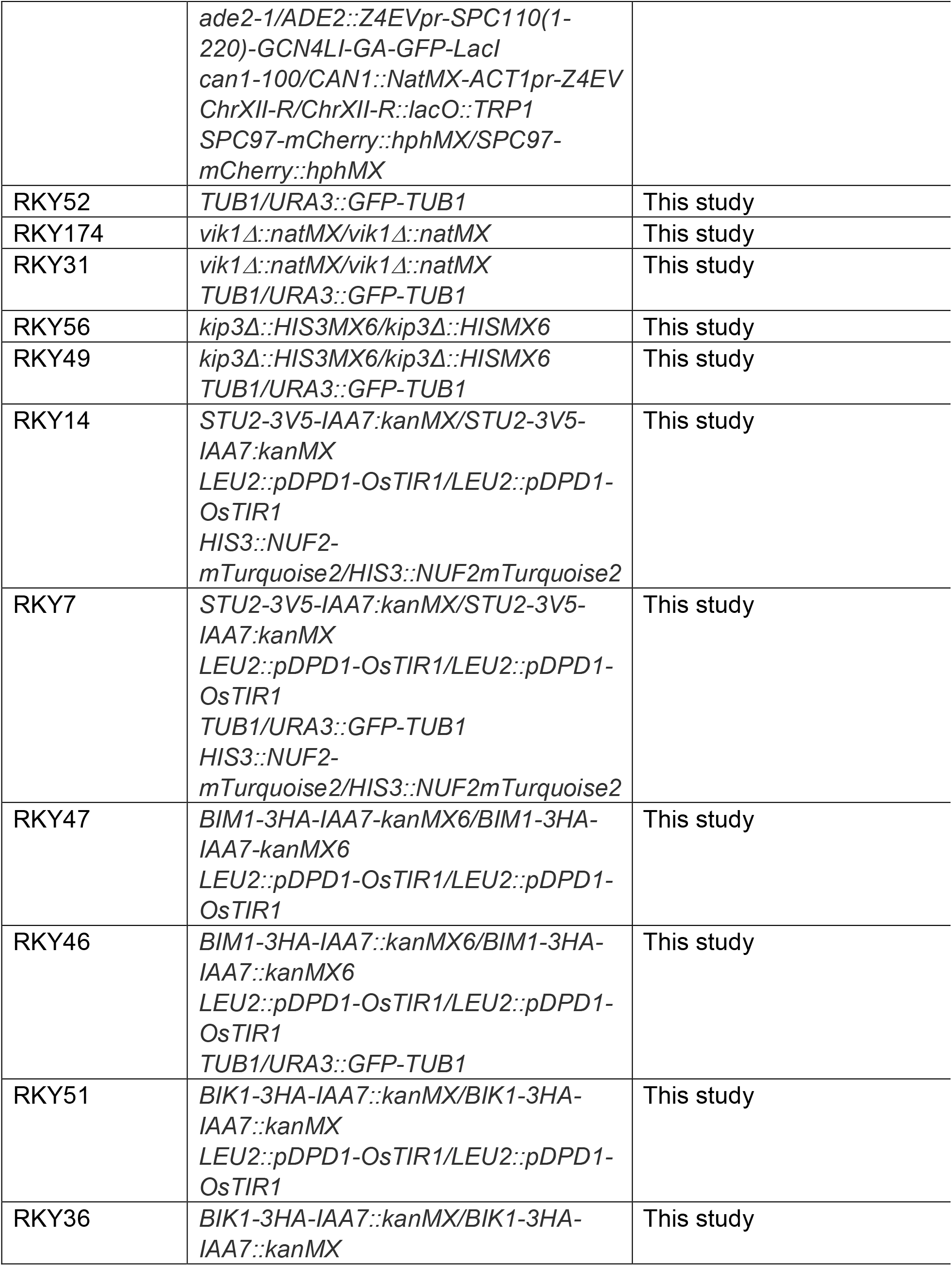
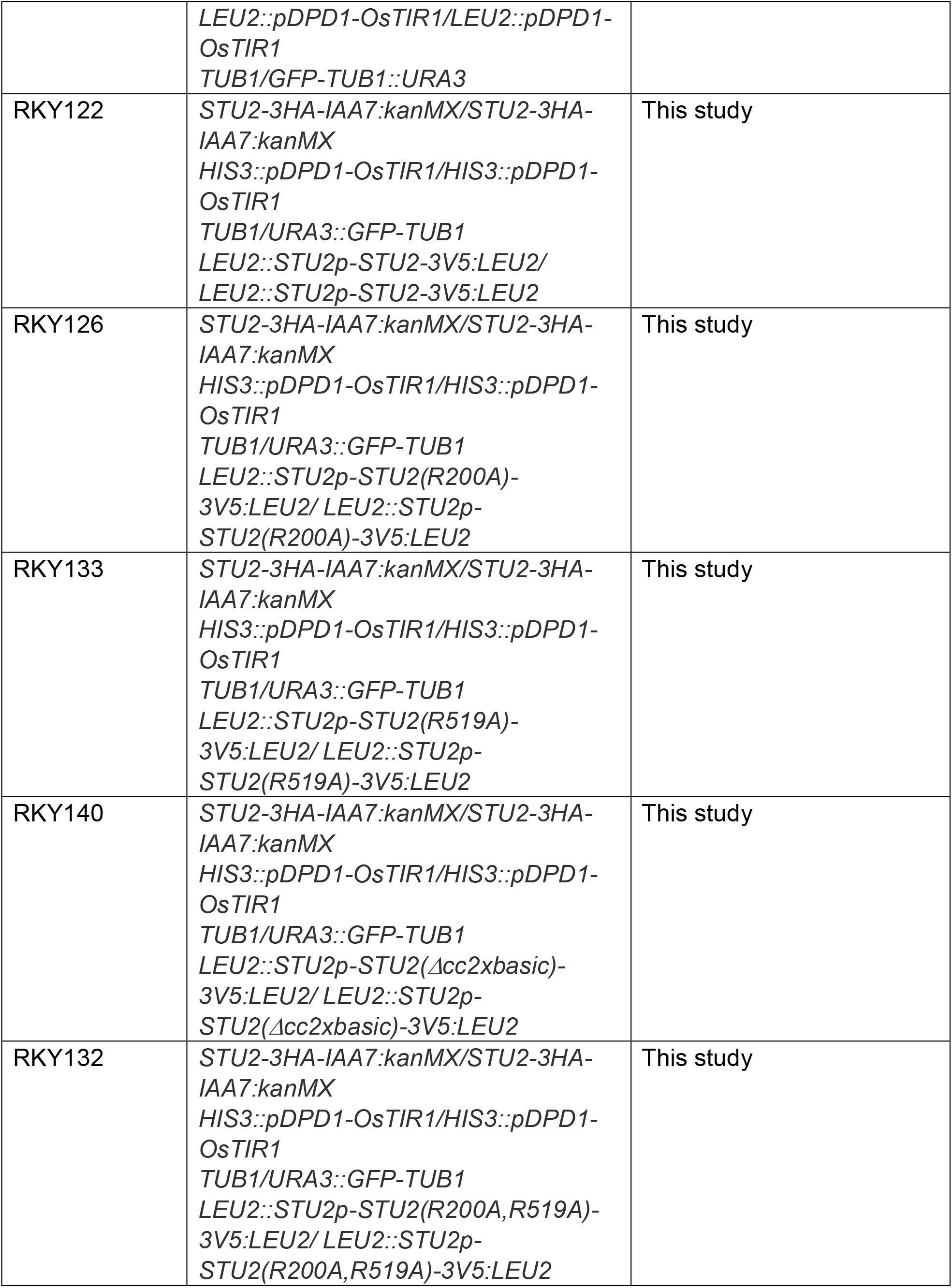
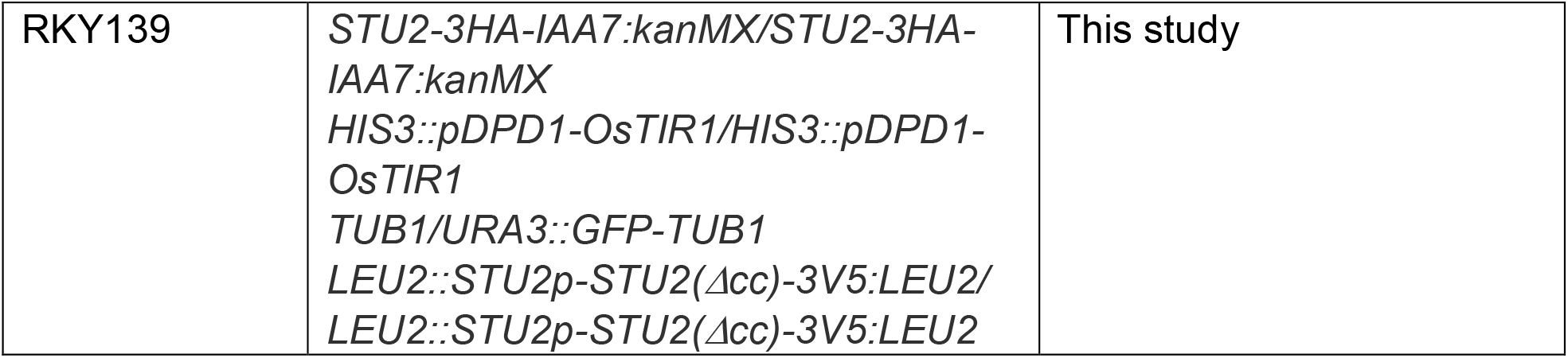
Strains used in this study^1^. ^1^ All strains contain the same markers as RKY44 except where noted.

| Strain name | Genotype | Established in |
| --- | --- | --- |
| TVY11-123D | *MATa his3-11,15 trp1-1 ADE2::Z4EVpr-SPC110(1-220)-GCN4LI-GA-GFP-LacI CAN1::NatMX-ACT1pr-Z4EV ChrXII-R::lacO::TRP1. SPC97-mCherry::hphMX LEU2::GFP-Tub1 SPC110-AID:kanMX URA3::OS-TIR1-myc3* | This study |
| RKY44 | *MATa/MATα his3-11,15/his3-11/15 leu2-3,112/leu2-3,112 trp1-1/trp1-1 ura3-1/ura3-1* | This study |
|  | *ade2-1/ADE2::Z4EVpr-SPC110(1-220)-GCN4LI-GA-GFP-LacI*<br>*can1-100/CAN1::NatMX-ACT1pr-Z4EV*<br>*ChrXII-R/ChrXII-R::lacO::TRP1*<br>*SPC97-mCherry::hphMX/SPC97-mCherry::hphMX* |  |
| RKY52 | *TUB1/URA3::GFP-TUB1* | This study |
| RKY174 | *vik1Δ::natMX/vik1Δ::natMX* | This study |
| RKY31 | *vik1Δ::natMX/vik1Δ::natMX*<br>*TUB1/URA3::GFP-TUB1* | This study |
| RKY56 | *kip3Δ::HIS3MX6/kip3Δ::HISMX6* | This study |
| RKY49 | *kip3Δ::HIS3MX6/kip3Δ::HISMX6*<br>*TUB1/URA3::GFP-TUB1* | This study |
| RKY14 | *STU2-3V5-IAA7:kanMX/STU2-3V5-IAA7:kanMX*<br>*LEU2::pDPD1-OsTIR1/LEU2::pDPD1-OsTIR1*<br>*HIS3::NUF2-mTurquoise2/HIS3::NUF2mTurquoise2* | This study |
| RKY7 | *STU2-3V5-IAA7:kanMX/STU2-3V5-IAA7:kanMX*<br>*LEU2::pDPD1-OsTIR1/LEU2::pDPD1-OsTIR1*<br>*TUB1/URA3::GFP-TUB1*<br>*HIS3::NUF2-mTurquoise2/HIS3::NUF2mTurquoise2* | This study |
| RKY47 | *BIM1-3HA-IAA7-kanMX6/BIM1-3HA-IAA7-kanMX6*<br>*LEU2::pDPD1-OsTIR1/LEU2::pDPD1-OsTIR1* | This study |
| RKY46 | *BIM1-3HA-IAA7::kanMX6/BIM1-3HA-IAA7::kanMX6*<br>*LEU2::pDPD1-OsTIR1/LEU2::pDPD1-OsTIR1*<br>*TUB1/URA3::GFP-TUB1* | This study |
| RKY51 | *BIK1-3HA-IAA7::kanMX/BIK1-3HA-IAA7::kanMX*<br>*LEU2::pDPD1-OsTIR1/LEU2::pDPD1-OsTIR1* | This study |
| RKY36 | *BIK1-3HA-IAA7::kanMX/BIK1-3HA-IAA7::kanMX* | This study |
|  | *LEU2::pDPD1-OsTIR1/LEU2::pDPD1-OsTIR1*<br>*TUB1/GFP-TUB1::URA3* |  |
| RKY122 | *STU2-3HA-IAA7:kanMX/STU2-3HA-IAA7:kanMX*<br>*HIS3::pDPD1-OsTIR1/HIS3::pDPD1-OsTIR1*<br>*TUB1/URA3::GFP-TUB1*<br>*LEU2::STU2p-STU2-3V5:LEU2/*<br>*LEU2::STU2p-STU2-3V5:LEU2* | This study |
| RKY126 | *STU2-3HA-IAA7:kanMX/STU2-3HA-IAA7:kanMX*<br>*HIS3::pDPD1-OsTIR1/HIS3::pDPD1-OsTIR1*<br>*TUB1/URA3::GFP-TUB1*<br>*LEU2::STU2p-STU2(R200A)-3V5:LEU2/*<br>*LEU2::STU2p-STU2(R200A)-3V5:LEU2* | This study |
| RKY133 | *STU2-3HA-IAA7:kanMX/STU2-3HA-IAA7:kanMX*<br>*HIS3::pDPD1-OsTIR1/HIS3::pDPD1-OsTIR1*<br>*TUB1/URA3::GFP-TUB1*<br>*LEU2::STU2p-STU2(R519A)-3V5:LEU2/*<br>*LEU2::STU2p-STU2(R519A)-3V5:LEU2* | This study |
| RKY140 | *STU2-3HA-IAA7:kanMX/STU2-3HA-IAA7:kanMX*<br>*HIS3::pDPD1-OsTIR1/HIS3::pDPD1-OsTIR1*<br>*TUB1/URA3::GFP-TUB1*<br>*LEU2::STU2p-STU2(Δcc2xbasic)-3V5:LEU2/*<br>*LEU2::STU2p-STU2(Δcc2xbasic)-3V5:LEU2* | This study |
| RKY132 | *STU2-3HA-IAA7:kanMX/STU2-3HA-IAA7:kanMX*<br>*HIS3::pDPD1-OsTIR1/HIS3::pDPD1-OsTIR1*<br>*TUB1/URA3::GFP-TUB1*<br>*LEU2::STU2p-STU2(R200A,R519A)-3V5:LEU2/*<br>*LEU2::STU2p-STU2(R200A,R519A)-3V5:LEU2* | This study |
| RKY139 | *STU2-3HA-IAA7:kanMX/STU2-3HA-IAA7:kanMX*<br>*HIS3::pDPD1-OsTIR1/HIS3::pDPD1-OsTIR1*<br>*TUB1/URA3::GFP-TUB1*<br>*LEU2::STU2p-STU2(Δcc)-3V5:LEU2/*<br>*LEU2::STU2p-STU2(Δcc)-3V5:LEU2* | This study |

**Table S2.** Plasmids used in this study.

| Plasmid name | Genetic markers | Established in |
| --- | --- | --- |
| pASF125 | For integration *TUB1-GFP-TUB1-URA3* at *URA3* | (Straight *et al.* 1997) |
| pGM25 | For integration pZ4EV-SPC110(1-220)-GCN4p1-GA-GFP-LacI-ADE2 at *ADE2* | (Lyon *et al.* 2016) |
| pGM22 | *LacOx256-TRP1* | (Lyon *et al.* 2016) |
| pSB2232 | For integration *STU2-3V5* at *LEU2* | (Miller *et al.* 2016) |
| pSB2306 | For integration *stu2(R200A)-3V5* at *LEU2* | (Miller *et al.* 2019) |
| pSB2307 | For integration *stu2(R519A)-3V5* at *LEU2* | (Miller *et al.* 2019) |
| pSB2308 | For integration of *stu2(R200A,R519A)-3V5* at *LEU2* | (Miller *et al.* 2019) |
| pSB2261 | For integration *stu2(Δ658-761::GDGAGL<sup>linker</sup>)* at *LEU2* | (Miller *et al.* 2019) |
| pSB3103 | For integration *stu2(2x_basicΔ658-761::GDGAGL<sup>linker</sup>)* at *LEU2* | (Miller *et al.* 2019) |

